# Spatial navigation impairment beyond episodic memory in autoimmune encephalitis

**DOI:** 10.64898/2026.08.24.746669

**Authors:** Sophia Rekers, Katharina Wurdack, Maron Mantwill, Antoine Coutrot, Guido Cammà, Joseph Kuchling, Harald Prüss, Michael Hornberger, Hugo Spiers, Carsten Finke

## Abstract

NMDAR and LGI1 encephalitis are the two most common forms of autoimmune encephalitis and are associated with persistent cognitive sequelae, particularly episodic memory impairment. Patients also report lasting difficulties with spatial orientation and navigation, yet these symptoms remain poorly characterized. Both disorders affect neural systems supporting spatial navigation, including prominent hippocampal pathology alongside cingulate, temporo-parietal, thalamic and cerebellar alterations identified in advanced neuroimaging studies. Here, we therefore investigated the frequency and clinical relevance of spatial navigation impairment in post-acute NMDAR and LGI1 encephalitis, its relationship with episodic memory dysfunction, and its structural correlates.

We included 80 post-acute patients from the autoimmune encephalitis outpatient clinic at Charité – Universitätsmedizin Berlin: 50 with NMDAR encephalitis (mean age 35.0 years, range 19–71; 90% female; median 6.9 years from onset) and 30 with LGI1 encephalitis (mean age 63.6 years, range 33–84; 67% male; median 2.7 years from onset). Spatial navigation was assessed using a passive map-assisted task (VIENNA Young) and an active wayfinding task (Sea Hero Quest), and its relationship with verbal episodic memory was examined using the Rey Auditory Verbal Learning Test. Structural MRI analyses assessed cortical thickness, subcortical volumes and diffusion measures in preselected navigation- and memory-related regions.

Patients with NMDAR and LGI1 encephalitis performed worse than matched controls on map-assisted navigation, and navigation performance showed strong convergence across the two navigation paradigms. Norm-referenced navigation impairment affected 57% of patients with NMDAR encephalitis and 70% with LGI1 encephalitis. In NMDAR encephalitis, selective navigation impairment was more common than selective memory impairment (41% versus 14%; χ² = 6.26, *p* = .012), supporting partial dissociation. In LGI1 encephalitis, navigation and memory impairments were similarly frequent and strongly overlapping, with 53% of patients impaired in both domains. Older age was a shared risk factor for navigation impairment. Structurally, NMDAR encephalitis showed partly distinct navigation- and memory-related alteration patterns, with navigation-specific parietal-paracentral and cerebellar abnormalities and memory-specific temporal-hippocampal-thalamic involvement. LGI1 encephalitis showed more widespread, predominantly memory-related alterations without a robust navigation-specific structural signature.

Our findings identify spatial navigation as a frequently affected but under-assessed cognitive domain in post-acute NMDAR and LGI1 encephalitis. They provide clinical evidence that navigation and episodic memory are partially dissociable yet overlapping functions whose degree of separability varies with the extent and distribution of network pathology. Incorporating norm-referenced navigation assessment into longitudinal follow-up could improve the characterization of cognitive profiles and related support needs, while reducing the risk that impairments relevant to everyday functioning and long-term quality of life remain undetected.

## Introduction

Anti-N-methyl-D-aspartate receptor (NMDAR) encephalitis^1^ and leucine-rich glioma-inactivated 1 (LGI1) encephalitis^2^ are the two most common variants of autoimmune encephalitis. Both variants particularly affect the hippocampus^3^ and are associated with long-term cognitive and neuropsychiatric sequelae^4–7^. Among these, persistent episodic memory impairment is the most consistently reported. In the post-acute phase, prevalence estimates for episodic memory impairments range from approximately 22-72% for NMDAR encephalitis and 20-75% for LGI1 encephalitis, depending on the assessment time point, neuropsychological instruments used and impairment thresholds^4,5,7–10^.

MRI studies have localized these deficits in both NMDAR and LGI1 encephalitis to a coherent limbic circuit, encompassing structural, microstructural and connectivity changes in the hippocampus and thalamus^9,11–13^, fornix^14,15^, cingulum^16–19^ and cerebellum^20–22^. This limbic circuit largely overlaps with the core network supporting spatial navigation, a critical function for everyday participation and safety^23–25^. Lesions to this network and adjacent entorhinal and posteromedial cortices (including the posterior cingulate cortex, retrosplenial cortex and precuneus) have consistently been linked to impairments in spatial navigation^26–34^, suggesting that they might represent a core clinical feature of autoimmune encephalitis. Indeed, more than half of post-acute patients with NMDAR and LGI1 encephalitis self-report problems with spatial orientation^6,35^ and a pilot study suggests that they might even exhibit impaired driving patterns^36^. However, while visuospatial memory and construction impairments have been identified using standard paper-and-pencil neuropsychological assessments^13,37^, these are not suited to capture more complex spatial navigation performance^38^, making it likely that challenges in this function remain undetected and thus untreated in many patients.

Here, we leverage recent advances in digital spatial navigation assessments to evaluate (i) whether post-acute NMDAR and LGI1 encephalitis patients exhibit navigation performance below matched controls and normative expectations; (ii) whether this navigation performance profile is dissociable from episodic memory performance; and (iii) how it relates to the structural integrity of limbic, cingulate, temporal and parietal regions. To this end, we employed a multi-modal approach using two validated, norm-referenced navigation tasks that tap into complementary cognitive processes, i.e. a passive map-assisted navigation assessment focusing on visuospatial/executive components (VIENNA Young)^39^ and an active wayfinding test that relies on spatial encoding and recall (Sea Hero Quest)^40^. We further examined the relationship between navigation performance and individual differences in macro- and microstructural integrity using high-resolution MRI and diffusion-tensor imaging. This integrative approach directly addresses key clinical and mechanistic gaps and provides a framework for incorporating spatial navigation testing into follow-up assessments of patients with autoimmune encephalitis.

## Materials and methods

### Recruitment and eligibility

We recruited 50 patients with NMDAR encephalitis and 30 patients with LGI1 encephalitis from the autoimmune encephalitis outpatient clinic at Charité – Universitätsmedizin Berlin. All patients fulfilled current diagnostic criteria and tested positive for NMDAR or LGI1 antibodies in serum and/or cerebrospinal fluid using indirect immunofluorescence assays. Only patients without concomitant herpes simplex virus encephalitis or neuroimmunological overlap syndromes, such as multiple sclerosis, were included.

Patients with NMDAR and LGI1 encephalitis were compared with separate age- and sex-matched healthy control groups. Clinical data were collected using standardized case report forms. The modified Rankin Scale (mRS) and the Clinical Assessment Scale in Autoimmune Encephalitis (CASE)^41^ were scored retrospectively by two clinically experienced investigators (GC and KW). All participants gave written informed consent and the study was approved by the local ethics committee of Charité – Universitätsmedizin Berlin (EA4/047/21 and EA4/011/19).

One patient was excluded from verbal episodic memory analyses because German was not their native language and one control lacked a verbal recognition score because the recognition trial was not administered. One patient from each cohort did not undergo MRI because of claustrophobia. One additional NMDAR encephalitis patient was excluded from imaging analyses because of an unrelated pre-existing structural brain abnormality; this patient performed within normal expectations on cognitive testing.

### Materials and procedure

Spatial navigation performance was evaluated using two paradigms with distinct and complementary degrees of memory involvement (see Fig. 1). The Virtual Environments Navigation Assessment for young and middle-aged adults (VIENNA Young^39^) is a map-assisted spatial navigation task comprising 12 virtual hallway scenes of increasing complexity. In each trial, participants watch a character move through a grid-like corridor while tracking the character’s position on a schematic map shown next to the video. When the video ends, they identify the door chosen by the character by clicking the corresponding door on the map. In VIENNA Young, spatial information is continuously available during each trial, so successful navigation does not rely on memorization. This paradigm thus emphasizes spatial and executive processing while minimizing demands on episodic memory consolidation and retrieval.

**Figure 1.**
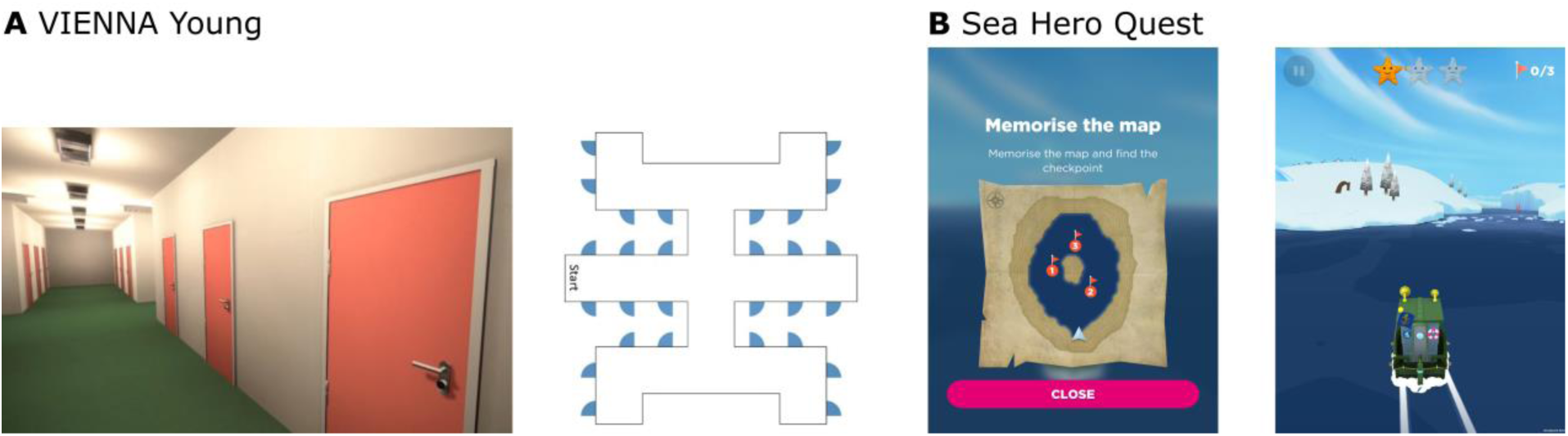
Visualization of the spatial navigation paradigms. (**A**) Virtual Environments Navigation Assessment for young and middle-aged adults (VIENNA Young; Level 7). Participants watch a first-person video of a character navigating a hallway environment while tracking their position on a schematic map that remains visible throughout the trial. At the end of each video, participants identify the door selected by the character. In later levels, the map is rotated to increase task difficulty. (**B**) Sea Hero Quest (Level 7). Participants first view a map indicating the starting position (arrowhead) and checkpoints (numbered circles). After the map is closed, participants actively steer a boat through the water environment to locate the checkpoints in the correct order.

In contrast, Sea Hero Quest (SHQ)^40^ is a video game in which participants actively navigate a boat through virtual water environments. Each wayfinding level used in this study consists of an initially presented map indicating the starting location and the location of several checkpoints that need to be memorized. After the map is closed by the participant, they steer the boat through the environment to locate the checkpoints in the specified order. We administered the practice levels 1 and 2 and the wayfinding levels 6, 7, 8, 11 and 21. Because SHQ was available only during part of the recruitment period, analyses were based on a smaller subset of patients. In the LGI1 encephalitis cohort, three participants did not complete levels 7 and 21. Their data from the remaining levels were retained because excluding participants whose non-completion may have reflected greater task difficulty could have biased performance estimates upward.

Verbal episodic memory performance was assessed using the German version of the Rey Auditory Verbal Learning Test (RAVLT), the Verbal Learning and Memory Test^42,43^. We examined three outcomes: verbal learning, defined as the total number of words correctly recalled across the five learning trials; delayed recall, defined as the number of words recalled after approximately 30 minutes; and word recognition, the number of correctly recognized words among targets and distractors.

### Norm-referenced scoring and classification

Spatial navigation and verbal episodic memory performance were evaluated against German normative data. In accordance with the respective norming recommendations, VIENNA Young and SHQ were compared to their respective age- and gender-matched reference groups, whereas RAVLT scores were normed in comparison to age-specific norms. The regression-based VIENNA Young normative models were developed for individuals aged up to 67 years. Since 15 patients (14 LGI1 encephalitis, 1 NMDAR encephalitis) were older than 67 years, we extrapolated the normative model to age 84; validation of this extrapolation is described in the Supplementary Methods. SHQ wayfinding performance was adjusted for dexterity using levels 1 and 2. Wayfinding and practice-level percentiles were logit-transformed. The mean-centred practice level percentile was then subtracted from the wayfinding percentile on the logit scale, and the adjusted score was inverse-logit transformed back to the 0–1 scale. For the RAVLT, when test scores corresponded to a percentile range, such as PR 50–75, the lower bound of the range was used for analyses.

For all tests, performance was classified according to American Academy of Clinical Neuropsychology guidelines^44^: below average (PR < 9), low average (9 ≤ PR ≤ 24) or within normal expectations (PR > 24). For binary analyses, below-average and low-average performance were combined and operationally defined as impaired performance (PR ≤ 24), in contrast to performance within normal expectations (PR > 24). To examine overlap between navigation and memory impairment, we additionally created domain-level classifications. The navigation domain comprised VIENNA Young and SHQ, whereas the verbal episodic memory domain comprised RAVLT learning, delayed recall and recognition. A domain was classified as unimpaired only when all constituent measures were within normal expectations and as impaired when at least one measure indicated performance below normal expectations.

### MRI data acquisition

All MRI data were acquired at the Berlin Centre for Advanced Neuroimaging using a 3 T MAGNETOM Prisma scanner (Siemens Healthineers, Erlangen, Germany). In the NMDAR encephalitis cohort, 43 patients were scanned using a 64-channel head coil and five using a 20- channel head coil; the corresponding numbers in the LGI1 encephalitis cohort were 17 and 12.

Structural imaging comprised a high-resolution 3D magnetisation-prepared rapid gradient-echo (MPRAGE) sequence with 1-mm isotropic voxels. The 64-channel protocol used a repetition time of 2500 ms, echo time of 2.64 ms, inversion time of 1000 ms and flip angle of 8°, whereas the 20- channel protocol used 1900 ms, 3.03 ms, 900 ms and 9°, respectively. For three participants, only a 0.8-mm isotropic acquisition was available and was used (echo time = 2.22 ms).

Diffusion analyses were restricted to the 64-channel protocol because the diffusion sequences differed substantially. The 64-channel protocol comprised two anterior-to-posterior phase-encoded runs with 1.5-mm isotropic voxels, 98 and 99 diffusion directions, b-values of 1500 and 3000 s/mm², a repetition time of 3230 ms, an echo time of 89.2 ms, monopolar diffusion encoding and a multiband acceleration factor of 4. Opposite phase-encoded spin-echo images were acquired for susceptibility-distortion correction.

### Imaging data analyses

Structural T1-weighted images were processed using FreeSurfer version 7.4.1 with the standard recon-all pipeline for automated subcortical segmentation and cortical surface reconstruction^45^. Cortical surfaces were parcellated according to the Desikan-Killiany atlas^46^. Cortical thickness in the parietal, temporal and cingulate cortices and bilateral hippocampal, thalamic and cerebellar grey matter volumes were extracted for analysis.

Diffusion-weighted images were preprocessed using QSIPrep version 1.0.2^47^, including MP-PCA denoising, Gibbs ringing removal, B1 bias field correction, eddy current, head motion and susceptibility distortion correction, and registration to the T1-weighted image. Diffusion tensors were fitted using DSI Studio (Hou version)^48^ as distributed in QSIRecon version 1.1.1 to derive fractional anisotropy and mean diffusivity maps. Subcortical regions of interest comprised the bilateral hippocampus, thalamus and cerebellar white matter. Masks were derived from the QSIPrep ACPC-space anatomical segmentation, eroded by one voxel to reduce partial-volume effects and resampled to the diffusion grid using nearest-neighbour interpolation before extraction of mean fractional anisotropy and mean diffusivity. White matter pathways, including the dorsal, peri-genual and temporal cingulum and the fornix, were reconstructed separately using XTRACT within FSL version 6.0^49,50^.

All structural and diffusion outputs underwent visual quality control. Segmentation, surface-reconstruction and tract-reconstruction errors were corrected where feasible; otherwise, affected measures were excluded. Regional imaging values were additionally screened within each group and region using MAD-based *z*-scores with |*z*| > 3.5 flagged for visual review. Values were excluded only when abnormalities reflected processing errors or focal pathology unrelated to autoimmune encephalitis; plausible anatomical variation and disease-related abnormalities were retained.

### Statistical analyses

Analyses were conducted in *R* (version 4.4.2)^51^, with α set at 0.05. Patients with NMDAR encephalitis and LGI1 encephalitis were matched separately 1:1 to healthy controls using optimal propensity-score matching on age and sex. Imaging pairs were additionally matched exactly on head-coil protocol. Paired analyses included complete patient–control pairs only. Prespecified directional comparisons were tested one-sided in the hypothesised direction; all other tests were two-sided. Where multiple regions were tested, *p*-values were adjusted using the Benjamini– Hochberg false discovery rate (FDR) procedure within each cohort, imaging modality and planned contrast.

Matched patient-control comparisons in navigation, memory and imaging measures were assessed using paired-sample *t*-tests, with Hedges’ *g* with small-sample correction reported as the effect size. Convergence between VIENNA Young and dexterity-adjusted SHQ performance was assessed using product-moment correlations. Differences between paired impairment classifications were assessed using McNemar tests, with exact tests used for sparse discordant pairs.

Candidate predictors of impaired navigation performance (PR ≤ 24) were examined separately in NMDAR and LGI1 encephalitis using binary logistic regression. Predictors comprised age, sex, years of education, years since disease onset, CASE impairment status at the study visit, highest CASE score during the acute disease phase, treatment delay, second-line therapy and rituximab treatment. Candidate models were compared using the corrected Akaike information criterion (AICc) on a common complete-case sample, and the best-supported model was refitted using the maximum available sample. Treatment variables were subsequently added individually and evaluated using likelihood-ratio tests and changes in model fit. Among NMDAR encephalitis patients with repeated assessments, first-to-last changes in VIENNA Young scores were evaluated using a one-sample Wilcoxon signed-rank test.

Potential effects of head-coil protocol on structural imaging measures were examined before pooling the data. Volumetric comparisons were adjusted for estimated total intracranial volume (eTIV). To identify imaging differences related to navigation or memory impairment, we compared impaired patients with their matched controls and with unimpaired patients. Comparisons between patient subgroups were adjusted for age, and volumetric models were additionally adjusted for eTIV. Regions that also differed between unimpaired patients and their matched controls were excluded from the impairment-focused results, as these findings were more likely to reflect general disease effects. The reported findings therefore comprise the remaining differences between impaired patients and controls, together with differences between impaired and unimpaired patients.

## Results

### Patient characteristics

We included 50 patients with NMDAR encephalitis (age: *M* = 35.0 [19–71] years; 90% female) and 30 patients with LGI1 encephalitis (age: *M* = 63.6 [33–84] years; 67% male). The median time from symptom onset to study visit was 6.9 years in NMDAR encephalitis (range: 0.3–18.9 years) and 2.7 years in LGI1 encephalitis (range: 0.3–10.9 years). Median CASE and mRS scores indicated no significant disability in NMDAR encephalitis (CASE: *M̃* = 0.0 [0–6]; mRS: *M̃* = 1.0 [0–3]) and mild disability in LGI1 encephalitis (CASE: *M̃* = 2.0 [0–6]; mRS: *M̃* = 2.0 [0–4]).

### Comparison with controls and convergence across navigation paradigms

We first assessed spatial navigation performance in both autoimmune encephalitis variants using VIENNA Young. Patients with NMDAR and LGI1 encephalitis showed significantly lower performance than their respective age- and sex-matched controls, as summarized in Table 1 and visualized in Supplementary Fig. 1. Furthermore, we identified a high convergence between map-assisted navigation (VIENNA Young score) and wayfinding navigation (dexterity-adjusted SHQ wayfinding distance) performance in the NMDAR encephalitis cohort (*r* = −0.60, *t*(27) = −3.90, *p* < .001) and the LGI1 encephalitis cohort (*r* = −0.58, *t*(10) = −2.23, *p =* .049).

**Table 1.**
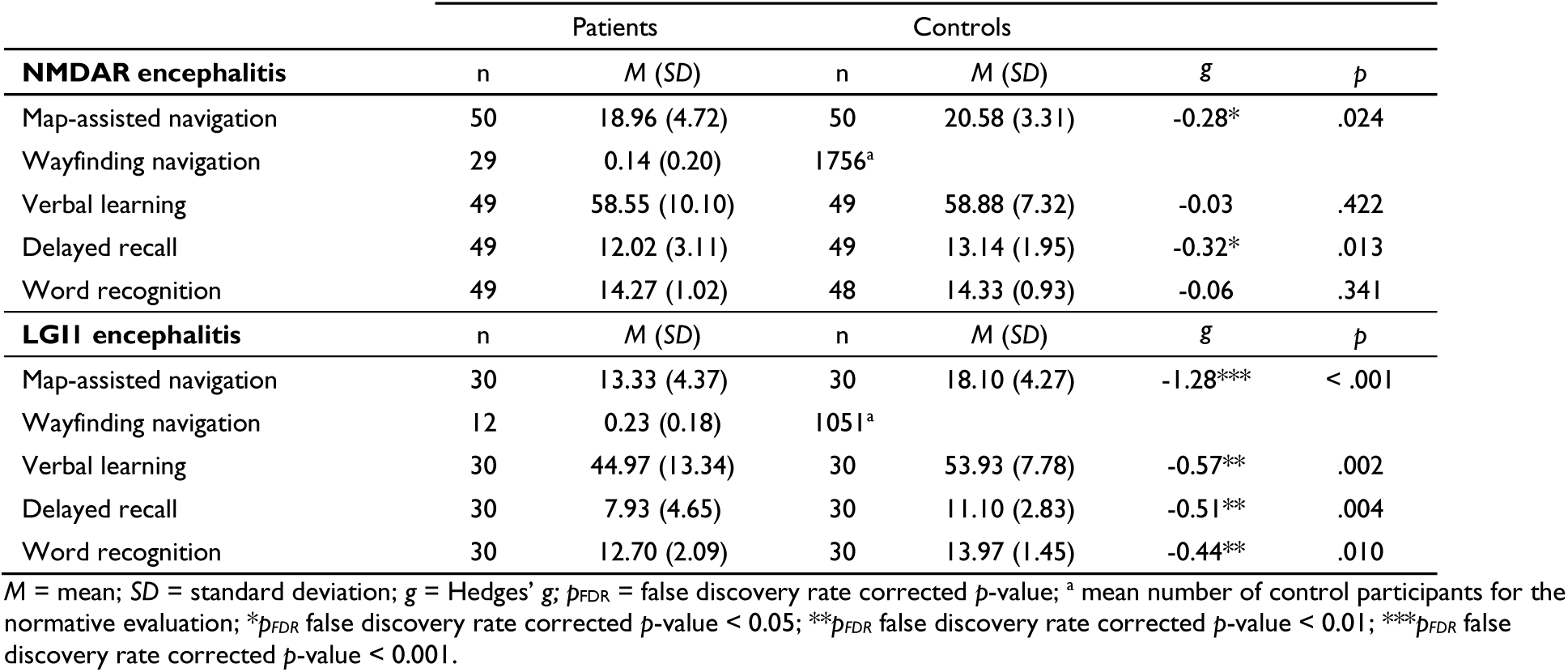
Descriptive statistics of the spatial navigation and verbal episodic memory performance.

|  | Patients |  | Controls |  |  |  |
| --- | --- | --- | --- | --- | --- | --- |
| **NMDAR encephalitis** | n | M (SD) | n | M (SD) | g | p |
| Map-assisted navigation | 50 | 18.96 (4.72) | 50 | 20.58 (3.31) | -0.28* | .024 |
| Wayfinding navigation | 29 | 0.14 (0.20) | 1756<sup>a</sup> |  |  |  |
| Verbal learning | 49 | 58.55 (10.10) | 49 | 58.88 (7.32) | -0.03 | .422 |
| Delayed recall | 49 | 12.02 (3.11) | 49 | 13.14 (1.95) | -0.32* | .013 |
| Word recognition | 49 | 14.27 (1.02) | 48 | 14.33 (0.93) | -0.06 | .341 |
| **LGII encephalitis** | n | M (SD) | n | M (SD) | g | p |
| Map-assisted navigation | 30 | 13.33 (4.37) | 30 | 18.10 (4.27) | -1.28*** | < .001 |
| Wayfinding navigation | 12 | 0.23 (0.18) | 1051<sup>a</sup> |  |  |  |
| Verbal learning | 30 | 44.97 (13.34) | 30 | 53.93 (7.78) | -0.57** | .002 |
| Delayed recall | 30 | 7.93 (4.65) | 30 | 11.10 (2.83) | -0.51** | .004 |
| Word recognition | 30 | 12.70 (2.09) | 30 | 13.97 (1.45) | -0.44** | .010 |

M = mean; SD = standard deviation; g = Hedges' g; $p_{FDR}$ = false discovery rate corrected p-value; <sup>a</sup> mean number of control participants for the normative evaluation; \* $p_{FDR}$ false discovery rate corrected p-value < 0.05; \*\* $p_{FDR}$ false discovery rate corrected p-value < 0.01; \*\*\* $p_{FDR}$ false discovery rate corrected p-value < 0.001.

### Overlap between navigation and memory performance

Next, we compared each cohort’s spatial navigation and verbal episodic memory performance with the respective normative data (see Fig. 2). In NMDAR encephalitis patients, map-assisted navigation performance was within normal expectations in 30 of 50 (60%) patients, 12 of 50 (24%) performed in the low-average range and 8 of 50 (16%) in the below-average range. A similar distribution was observed for wayfinding navigation, with 16 of 29 patients (55%) performing within normal expectations, 7 of 29 (24%) in the low-average range and 6 of 29 (21%) below average. By contrast, verbal episodic memory performance was largely preserved, with normal performance in 42 of 49 (86%) patients for learning, 42 of 49 (86%) for delayed recall and 39 of 49 (80%) for recognition. Low-average performance was observed in 4/49 (8%) for learning, 0/49 (0%) for delayed recall and 8/49 (16%) for recognition. Below-average performance was observed in 3/49 (6%), 7/49 (14%) and 2/49 (4%), respectively.

**Figure 2.**
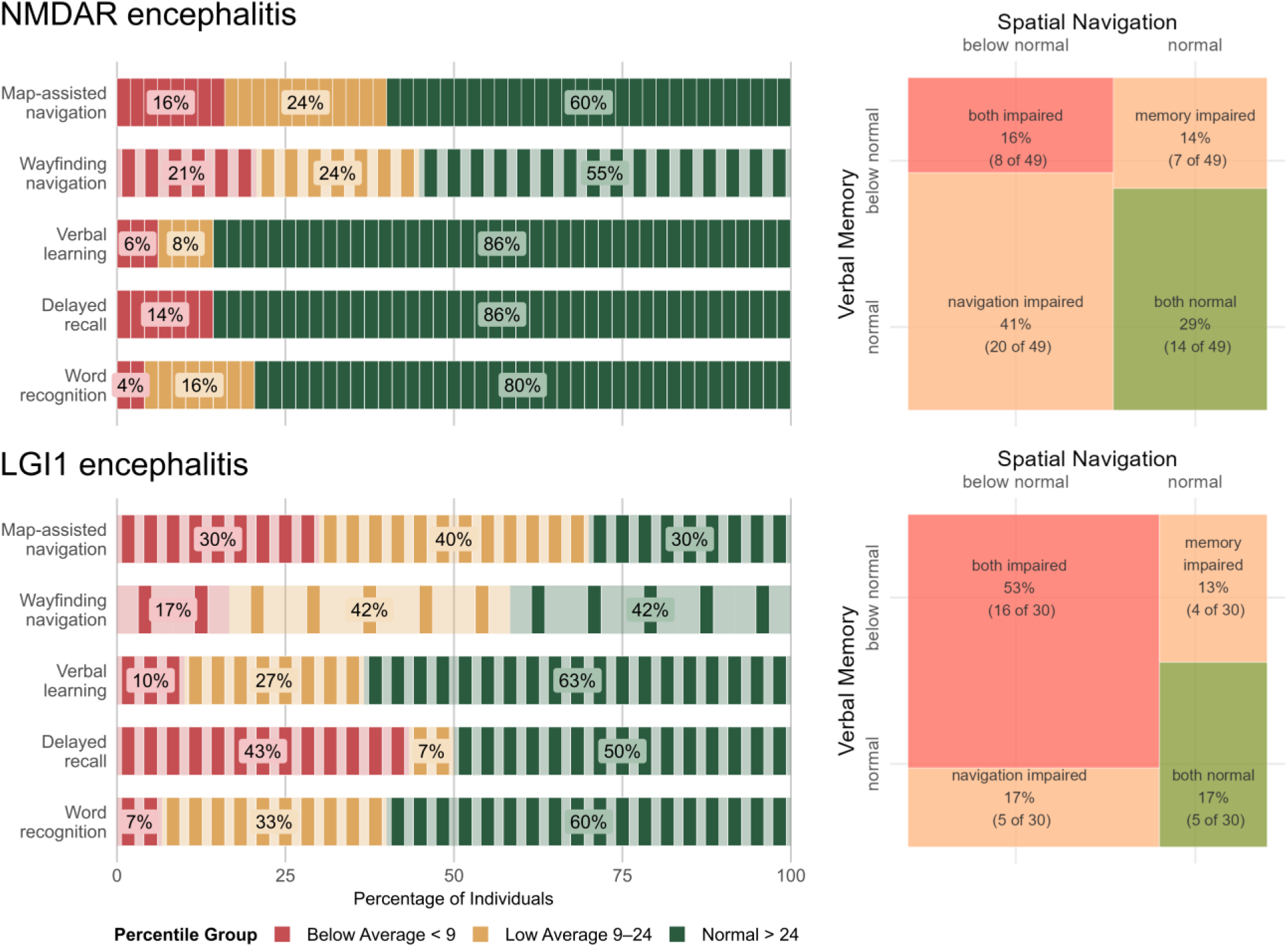
Distribution and overlap of spatial navigation and episodic memory performance. Left, participant- level tile charts showing the distribution of performance categories across map-assisted navigation measured with VIENNA Young, wayfinding measured with Sea Hero Quest (SHQ) and Rey Auditory Verbal Learning Test (RAVLT) learning, delayed recall and recognition in NMDAR encephalitis (top) and LGI1 encephalitis (bottom). Each rectangle represents one participant and rows are scaled to 100%. Colours correspond to the percentile categories shown in the figure key. Right, mosaic plots show the overlap between binary navigation and memory classifications. Rectangle areas are proportional to the number of patients in each category. Navigation (VIENNA Young and SHQ) and memory (RAVLT learning, delayed recall and recognition) were classified as impaired when any available measure in the domain was below normal expectations (percentile rank, PR ≤ 24), and as normal when all available measures were within normal expectations (PR > 24).

In the LGI1 cohort, memory and navigation were more frequently impaired. Map-assisted navigation was within normal expectations in 9/30 (30%) patients, low average in 12/30 (40%) and below-average in 9/30 (30%). Wayfinding navigation showed a similar distribution, with 5 of 12 patients (42%) performing within normal expectations, 5 of 12 (42%) in the low-average range and 2 of 12 (17%) below average. For verbal episodic memory, normal performance was observed in 19/30 (63%) patients for learning, 15/30 (50%) for delayed recall, and 18/30 (60%) for recognition. Low-average performance was observed in 8/30 (27%), 2/30 (7%), and 10/30 (33%), respectively; below-average performance was observed in 3/30 (10%), 13/30 (43%), and 2/30 (7%).

Binary categorization at PR ≤ 24 showed that, among 49 NMDAR encephalitis patients with available navigation and verbal memory classifications, 8 of 49 (16%) performed below normal in both domains, whereas 20 of 49 (41%) showed selective navigation impairment, 7 of 49 (14%) showed selective memory impairment, and 14 of 49 (29%) performed within normal expectations in both domains. Selective navigation impairment was significantly more frequent than selective memory impairment (*χ*^2^(1) = 6.26, *p* = .012). Among 30 LGI1 encephalitis patients, 16 of 30 (53%) performed below normal in both domains, 5 of 30 (17%) showed navigation impairment only, 4 of 30 (13%) showed memory impairment only, and 5 of 30 (17%) performed within normal expectations in both domains. The frequencies of isolated navigation and memory impairment did not differ (exact McNemar test, *p* > .999).

Overall, below-normal navigation performance was observed in 28 of 49 (57%) NMDAR encephalitis patients and 21 of 30 (70%) LGI1 encephalitis patients. Below-normal verbal memory performance was observed in 15 of 49 (31%) NMDAR and 20 of 30 (67%) LGI1 encephalitis patients. A relative predominance of navigation impairment over memory impairment was also evident in both cohorts when verbal memory was replaced with visual memory measures and was not explained by executive-function impairment (Supplementary Results). In supplementary analyses using continuous test scores to account for differences between normative references, working memory, but neither delayed recall nor attention, was associated with navigation performance in NMDAR encephalitis after adjustment for age, sex and education (Supplementary Results). None of the cognitive measures was significantly associated with navigation performance in LGI1 encephalitis.

### Risk factors for navigation impairment

Next, we explored candidate predictors of below-normal navigation performance. In the NMDAR encephalitis cohort, the best-supported model included age and time since disease onset (AICc = 61.57). Each year above the mean age was associated with a 5% increase in the odds of below-normal navigation performance (OR = 1.05, 95% CI [0.99, 1.11], *p* = .076), while each additional year since disease onset was associated with a 16% increase in the odds (OR = 1.16, 95% CI [1.01, 1.33], *p* = .036). The model was not significantly improved by adding treatment delay (*χ*^2^(1) = 0.27, *p* = .601, *Δ*AICc = 2.11), second-line therapy (*χ*^2^(1) = 0.17, *p* = .682, *Δ*AICc = 2.22), or specifically rituximab treatment (*χ*^2^(1) = 0.59, *p* = .443, *Δ*AICc = 1.79). To examine whether the cross-sectional association with time since onset was also reflected in within-person change, we conducted an exploratory follow-up analysis in 20 NMDAR encephalitis patients with navigation data from multiple visits. Individual VIENNA Young score trajectories were heterogeneous, with no consistent improvement or decline over time. The median within-person change was 0.0 points (*p* > .999).

In the LGI1 encephalitis cohort, the model with the lowest AICc included only age (AICc = 29.48). Each year above the mean age was associated with a 13% increase in the odds of below-normal navigation performance (OR = 1.13, 95% CI [1.03, 1.24], *p* = .011). The model was not significantly improved by adding treatment delay (*χ*^2^(1) = 0.31, *p* = .579, *Δ*AICc = 2.21), second-line therapy (*χ*^2^(1) = 0.24, *p* = .627, *Δ*AICc = 2.31) or specifically rituximab treatment (*χ*^2^(1) = 0.02, *p* = .894, *Δ*AICc = 2.46).

### Association with macrostructural imaging markers

We first assessed macrostructural differences between the autoimmune encephalitis cohorts and their matched controls in parietal, temporal and cingulate cortical thickness and in eTIV-adjusted volumes of the bilateral hippocampus, thalamus, and cerebellar grey matter. In the NMDAR encephalitis cohort (Table S1), cortical thickness was reduced in the bilateral superior temporal, left middle temporal, paracentral and caudal anterior cingulate regions, as well as in the right inferior parietal cortex. However, none of these differences survived correction for multiple comparisons (FDR). Subcortical analyses showed smaller right hippocampal and bilateral cerebellar grey matter volumes, which remained significant after FDR correction.

In LGI1 encephalitis patients (Table S2), cortical thickness was reduced across widespread temporo-parietal, medial parietal, and cingulate regions. After FDR correction, differences remained significant bilaterally in the fusiform, inferior temporal, supramarginal, precuneus and isthmus cingulate cortices, in the left entorhinal, middle temporal and inferior parietal regions, as well as in the right posterior cingulate cortex. Additional thinning in the bilateral banks STS (superior temporal sulcus), paracentral and superior parietal cortices, as well as right superior temporal, postcentral and inferior parietal cortices and smaller bilateral hippocampal volumes did not survive FDR correction.

We next examined structural differences specifically associated with navigation or memory impairment (Fig. 3). Patients with below-normal performance were compared with their matched healthy controls and with patients showing normal performance. Patient subgroup comparisons were adjusted for age in cortical thickness analyses and for age and eTIV in volume analyses. Only regions showing nominally significant differences are included in Fig. 3 and Tables 2 and 3.

**Figure 3.**
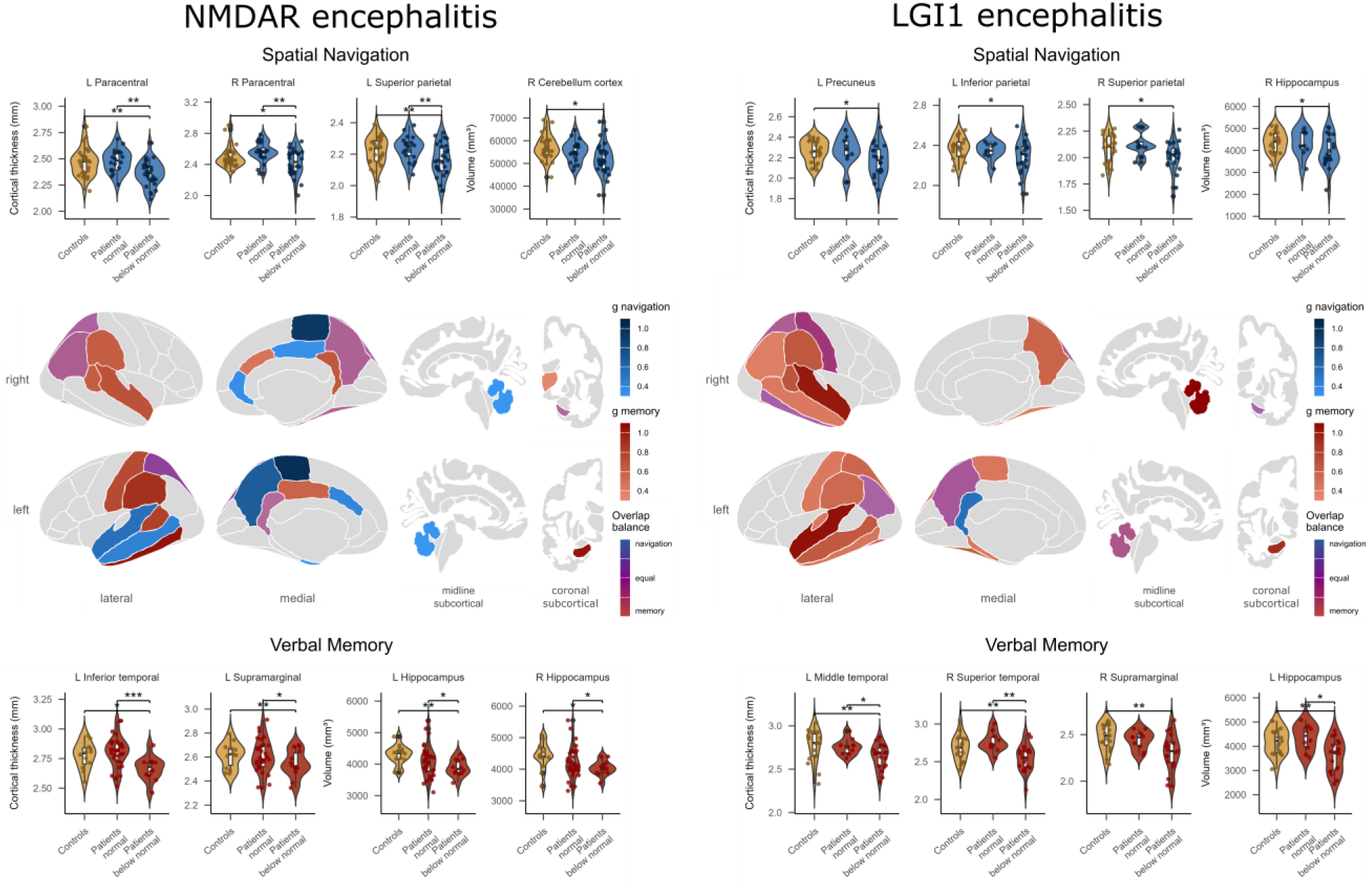
Distinct and overlapping macrostructural correlates of spatial navigation and episodic memory impairment. Cortical thickness and subcortical volume differences are shown separately for the NMDAR encephalitis cohort (left) and LGI1 encephalitis cohort (right), with navigation-related findings in the upper panels and memory-related findings in the lower panels. Brain maps show regions with nominally significant reductions in patients with below-normal performance relative to matched healthy controls or patients with normal performance. Colours indicate findings observed only in the navigation analyses (blue), only in the memory analyses (red) or in both domains (purple). For domain-specific regions, colour intensity reflects effect size; for overlapping regions, colour balance reflects the relative magnitude of the navigation- and memory-related effect sizes. Violin plots illustrate selected regional findings across matched controls, patients with normal performance, and patients with below-normal performance. Horizontal brackets indicate significant group comparisons, with * p < .05, ** p < .01 and *** p < .001.

**Table 2.** Significant macrostructural differences by performance status in the NMDAR encephalitis cohort.

| Navigation | Left hemisphere |  |  |  |  | Right hemisphere |  |  |  |  |
| --- | --- | --- | --- | --- | --- | --- | --- | --- | --- | --- |
|  | < normal | controls | normal | *g* | *p* | < normal | controls | normal | *g* | *p* |
| **Mean cortical thickness (mm)** |  |  |  |  |  |  |  |  |  |  |
| Entorhinal | 3.14 | 3.25 |  | -0.31 | .050 |  |  |  |  |  |
| Fusiform |  |  |  |  |  | 2.67 | 2.73 |  | -0.32 | .050 |
| Superior temporal | 2.74 | 2.87 |  | -0.54 | .003 |  |  |  |  |  |
|  | 2.76 |  | 2.84 | -0.54 | .033 |  |  |  |  |  |
| Middle temporal | 2.79 | 2.87 |  | -0.42 | .014 |  |  |  |  |  |
| Paracentral | 2.36 | 2.46 |  | -0.57 | .002 | 2.42 | 2.50 |  | -0.34 | .037 |
|  | 2.36 |  | 2.48 | -1.00* | .001 | 2.42 |  | 2.56 | -0.94* | .001 |
| Superior parietal | 2.16 | 2.22 |  | -0.50 | .006 | 2.16 | 2.22 |  | -0.40 | .019 |
|  | 2.16 |  | 2.24 | -0.85* | .004 |  |  |  |  |  |
| Inferior parietal |  |  |  |  |  | 2.50 | 2.56 |  | -0.39 | .022 |
| Precuneus | 2.36 | 2.41 |  | -0.41 | .017 | 2.38 | 2.43 |  | -0.36 | .031 |
|  | 2.36 |  | 2.42 | -0.68 | .021 |  |  |  |  |  |
| Isthmus-cingulate | 2.31 | 2.39 |  | -0.33 | .043 |  |  |  |  |  |
| Posterior-cingulate |  |  |  |  |  | 2.36 | 2.44 |  | -0.34 | .036 |
| Caudal anterior cingulate | 2.47 | 2.60 |  | -0.41 | .018 |  |  |  |  |  |
| Rostral anterior cingulate |  |  |  |  |  | 2.61 | 2.68 |  | -0.35 | .035 |
| **Mean eTIV adjusted volume (mm<sup>3</sup>)** |  |  |  |  |  |  |  |  |  |  |
| Hippocampus |  |  |  |  |  | 4,115 | 4,414 |  | -0.41 | .038 |
| Cerebellum | 51,747 | 56,412 |  | -0.31 | .025 | 52,184 | 57,044 |  | -0.31 | .023 |
| **Memory** |  |  |  |  |  |  |  |  |  |  |
| **Mean cortical thickness (mm)** |  |  |  |  |  |  |  |  |  |  |
| Fusiform |  |  |  |  |  | 2.61 |  | 2.72 | -0.83 | .011 |
| Superior temporal |  |  |  |  |  | 2.71 |  | 2.82 | -0.69 | .028 |
| Inferior temporal | 2.67 | 2.77 |  | -0.73 | .010 |  |  |  |  |  |
|  | 2.67 |  | 2.80 | -1.04* | < .001 |  |  |  |  |  |
| Banks STS | 2.36 | 2.49 |  | -0.60 | .017 |  |  |  |  |  |
|  | 2.35 |  | 2.48 | -0.80 | .010 | 2.50 |  | 2.61 | -0.62 | .036 |
| Postcentral | 2.06 | 2.13 |  | -0.52 | .030 |  |  |  |  |  |
|  | 2.06 |  | 2.13 | -0.74 | .014 |  |  |  |  |  |
| Supramarginal | 2.54 | 2.62 |  | -0.90 | .002 |  |  |  |  |  |
|  | 2.54 |  | 2.60 | -0.50 | .042 | 2.54 |  | 2.61 | -0.62 | .036 |
| Superior parietal | 2.15 |  | 2.21 | -0.67 | .034 | 2.13 |  | 2.20 | -0.63 | .025 |
| Inferior parietal |  |  |  |  |  | 2.46 | 2.54 |  | -0.79 | .004 |
|  |  |  |  |  |  | 2.46 |  | 2.54 | -0.75 | .013 |
| Precuneus |  |  |  |  |  | 2.35 |  | 2.44 | -0.72 | .006 |
| Isthmus-cingulate |  |  |  |  |  | 2.23 | 2.34 |  | -0.57 | .021 |
|  | 2.23 |  | 2.34 | -0.60 | .016 | 2.23 |  | 2.33 | -0.66 | .018 |
| Posterior-cingulate | 2.34 |  | 2.43 | -0.58 | .049 |  |  |  |  |  |
| Caudal anterior cingulate |  |  |  |  |  | 2.28 | 2.38 |  | -0.46 | .045 |
| **Mean eTIV adjusted volume (mm<sup>3</sup>)** |  |  |  |  |  |  |  |  |  |  |
| Hippocampus | 3,885 | 4,291 |  | -1.32** | .001 | 4,031 | 4,416 |  | -0.75 | .021 |
|  | 3,881 |  | 4,086 | -0.45 | .015 | 4,018 |  | 4,204 | -0.42 | .034 |
| Thalamus |  |  |  |  |  | 7,070 |  | 7,375 | -0.36 | .038 |

*g* = Hedges' *g*; \**p*<sub>FDR</sub> false discovery rate corrected *p*-value < 0.05; \*\**p*<sub>FDR</sub> false discovery rate corrected *p*-value < 0.01.

**Table 3.** Significant macrostructural differences by performance status in the LGI1 encephalitis cohort.

| Navigation | Left hemisphere |  |  |  |  | Right hemisphere |  |  |  |  |
| --- | --- | --- | --- | --- | --- | --- | --- | --- | --- | --- |
|  | < normal | controls | normal | g | p | < normal | controls | normal | g | p |
| **Mean cortical thickness (mm)** |  |  |  |  |  |  |  |  |  |  |
| Inferior temporal |  |  |  |  |  | 2.63 | 2.70 |  | -0.46 | .023 |
| Postcentral |  |  |  |  |  | 1.90 | 2.00 |  | -0.58 | .007 |
| Superior parietal |  |  |  |  |  | 2.00 | 2.08 |  | -0.40 | .038 |
| Inferior parietal | 2.29 | 2.37 |  | -0.42 | .032 |  |  |  |  |  |
| Precuneus | 2.17 | 2.27 |  | -0.49 | .016 |  |  |  |  |  |
| Isthmus-cingulate | 2.20 | 2.29 |  | -0.53 | .011 |  |  |  |  |  |
| **Mean eTIV adjusted volume (mm<sup>3</sup>)** |  |  |  |  |  |  |  |  |  |  |
| Hippocampus |  |  |  |  |  | 3,945 | 4,369 |  | -0.44 | .025 |
| Cerebellum | 51,345 | 55,470 |  | -0.40 | .012 |  |  |  |  |  |
| **Memory** |  |  |  |  |  |  |  |  |  |  |
| **Mean cortical thickness (mm)** |  |  |  |  |  |  |  |  |  |  |
| Parahippocampal | 2.52 | 2.74 |  | -0.44 | .031 |  |  |  |  |  |
| Fusiform | 2.55 | 2.66 |  | -0.61* | .006 | 2.54 | 2.65 |  | -0.41 | .038 |
| Superior temporal |  |  |  |  |  | 2.55 | 2.69 |  | -0.61* | .006 |
|  | 2.60 |  | 2.74 | -1.05 | .008 | 2.57 |  | 2.72 | -1.02 | .009 |
| Middle temporal | 2.64 | 2.77 |  | -0.65* | .004 | 2.65 | 2.77 |  | -0.43 | .034 |
|  | 2.66 |  | 2.73 | -0.55 | .036 |  |  |  |  |  |
| Inferior temporal | 2.60 | 2.71 |  | -0.46 | .026 | 2.62 | 2.70 |  | -0.46 | .026 |
| Transverse temporal | 2.08 | 2.22 |  | -0.44 | .030 |  |  |  |  |  |
| Banks STS |  |  |  |  |  | 2.37 | 2.52 |  | -0.68* | .003 |
|  |  |  |  |  |  | 2.38 |  | 2.49 | -0.65 | .040 |
| Paracentral | 2.28 | 2.37 |  | -0.45 | .029 |  |  |  |  |  |
| Postcentral | 1.93 | 2.01 |  | -0.43 | .034 | 1.87 | 2.02 |  | -0.91* | < .001 |
|  |  |  |  |  |  | 1.89 |  | 2.04 | -1.21** | < .001 |
| Supramarginal | 2.34 | 2.46 |  | -0.54* | .012 | 2.31 | 2.47 |  | -0.66* | .004 |
| Superior parietal | 2.05 | 2.13 |  | -0.40 | .044 | 1.99 | 2.10 |  | -0.54* | .012 |
|  | 2.06 |  | 2.13 | -0.56 | .044 | 2.00 |  | 2.10 | -0.77 | .030 |
| Inferior parietal | 2.28 | 2.37 |  | -0.51 | .016 | 2.30 | 2.38 |  | -0.40 | .044 |
| Precuneus | 2.15 | 2.28 |  | -0.66* | .004 | 2.17 | 2.31 |  | -0.57* | .009 |
| **Mean eTIV adjusted volume (mm<sup>3</sup>)** |  |  |  |  |  |  |  |  |  |  |
| Hippocampus | 3,590 | 4,197 |  | -0.58* | .009 | 3,889 | 4,377 |  | -0.45* | .020 |
|  | 3,652 |  | 4,238 | -0.91* | .013 |  |  |  |  |  |
| Cerebellum | 50,855 | 55,359 |  | -0.45* | .023 | 52,162 | 56,532 |  | -0.38 | .047 |
|  | 51,478 |  | 55,992 | -0.92 | .033 | 52,500 |  | 58,440 | -1.21* | .007 |

g = Hedges' g; \* $p_{FDR}$ false discovery rate corrected $p$ -value < 0.05; \*\* $p_{FDR}$ false discovery rate corrected $p$ -value < 0.01.

In the NMDAR encephalitis cohort, reductions showed partly distinct patterns for navigation and memory (details are provided in Table 2). Navigation-specific reductions involved left entorhinal and lateral temporal cortex; bilateral paracentral and left precuneus cortices; bilateral anterior and right posterior cingulate cortices; and bilateral cerebellar volumes. Reductions overlapping with the memory analyses involved the right fusiform cortex; bilateral superior, right inferior parietal and precuneus cortices; left isthmus cingulate cortex; and right hippocampal volume. Memory-specific reductions involved bilateral lateral temporal and supramarginal cortices; left postcentral cortex; left posterior and right isthmus and caudal anterior cingulate cortices; and left hippocampal and right thalamic volumes. After FDR correction, navigation-specific reductions remained significant in the bilateral paracentral cortex and left superior parietal cortex, and memory-specific reductions remained significant in the left inferior temporal cortex and left hippocampal volume.

In the LGI1 encephalitis cohort (Table 3), navigation-specific reductions were limited to the left isthmus cingulate cortex. Reductions overlapping with the memory analyses involved the right inferior temporal cortex; the left inferior parietal and precuneus and right postcentral and superior parietal cortices; and right hippocampal and left cerebellar volumes. Memory-specific reductions involved the bilateral ventral and lateral and left medial temporal cortices; the bilateral lateral parietal, left paracentral and precuneus cortices; and left hippocampal and right cerebellar volumes. After FDR correction, no navigation-specific reductions remained significant, whereas memory-specific reductions remained significant in lateral temporal cortices in both hemispheres, with additional left ventral temporal involvement; bilateral lateral and medial parietal cortices; and bilateral hippocampal and cerebellar volumes.

Across both autoimmune encephalitis variants, convergent navigation-specific reductions involved the left precuneus, left isthmus cingulate cortex and left cerebellar volume. Convergent reductions shared between navigation and memory analyses involved the right superior parietal cortex and right hippocampal volume. Convergent memory-specific reductions involved the left inferior temporal cortex and the right superior temporal, fusiform, and banks STS cortices; the bilateral supramarginal cortices, left superior parietal and postcentral cortices, and right inferior parietal and precuneus cortices; as well as lower left hippocampal volume.

### Association with microstructural imaging markers

We then assessed group differences in mean diffusivity (MD) and fractional anisotropy (FA) between each autoimmune encephalitis cohort and its matched control group. NMDAR encephalitis patients showed higher bilateral cerebellar white matter MD and lower bilateral hippocampal FA compared to matched controls, with all four differences surviving FDR correction (Table S3). Nominal differences additionally included higher left hippocampal and bilateral temporal cingulum MD and lower bilateral cerebellar white matter FA. LGI1 encephalitis patients showed higher bilateral hippocampal MD and lower bilateral hippocampal FA (Table S4). Higher left temporal cingulum MD and lower right temporal cingulum FA were also observed. However, none of these differences survived FDR correction.

We next examined microstructural differences specifically associated with navigation or memory impairment, analogous to the previous approach of the macrostructural associations. In NMDAR encephalitis (Table 4), navigation-specific findings comprised higher MD in the right cerebellar white matter and lower FA in the bilateral hippocampi, right cerebellar white matter, and right fornix. Higher right cerebellar MD and lower left hippocampal FA survived FDR correction. Higher left temporal cingulum MD was observed in both navigation and memory analyses, although only the memory-related difference survived FDR correction. Memory-specific analyses comprised higher MD in the left hippocampus and right temporal cingulum and lower FA in the bilateral thalami. Higher left hippocampal MD and lower bilateral thalamic FA survived FDR correction.

**Table 4.** Significant microstructural differences by performance status in the NMDAR encephalitis cohort.

| Navigation | DTI metric | Left hemisphere |  |  |  |  |  | Right hemisphere |  |  |  |  |  |
| --- | --- | --- | --- | --- | --- | --- | --- | --- | --- | --- | --- | --- | --- |
|  |  | n | < normal | controls | normal | g | p | n | < normal | controls | normal | g | p |
| Hippocampus | FA | 25 | 0.117 | 0.130 |  | -0.67* | .001 | 25 | 0.118 | 0.127 |  | -0.50 | .008 |
|  | FA |  |  |  |  |  |  | 25/16 | 0.120 |  | 0.130 | -0.89 | .017 |
| Cerebellar white matter | MD |  |  |  |  |  |  | 26 | 0.603 | 0.590 |  | 0.71** | < .001 |
|  | FA |  |  |  |  |  |  | 26 | 0.454 | 0.467 |  | -0.36 | .034 |
| Temporal cingulum | MD | 25 | 0.750 | 0.738 |  | 0.38 | .032 |  |  |  |  |  |  |
| Fornix | FA |  |  |  |  |  |  | 13/4 | 0.253 |  | 0.366 | -2.20 | .005 |
| **Memory** |  |  |  |  |  |  |  |  |  |  |  |  |  |
| Hippocampus | MD | 10 | 0.810 | 0.778 |  | 0.73* | .017 |  |  |  |  |  |  |
| Thalamus | FA | 12 | 0.285 | 0.295 |  | -0.72* | .011 | 12 | 0.280 | 0.289 |  | -1.48** | < .001 |
| Temporal cingulum | MD | 11 | 0.760 | 0.736 |  | 0.69* | .016 | 11 | 0.756 | 0.736 |  | 0.57 | .034 |

DTI = diffusion tensor imaging; FA = fractional anisotropy; MD = mean diffusivity; g = Hedges' g; MD values are reported in $10^{-3}$ mm<sup>2</sup>/s. \* $p_{FDR}$ false discovery rate corrected $p$ -value < 0.05; \*\* $p_{FDR}$ false discovery rate corrected $p$ -value < 0.01.

In LGI1 encephalitis (Table 5), lower right hippocampal FA was navigation-specific, while higher right hippocampal MD was observed in both navigation and memory analyses. Memory-specific findings comprised higher MD in the left hippocampus, right cerebellar white matter, and left temporal cingulum and lower FA in the left hippocampus and right temporal cingulum. None of the domain-specific LGI1 encephalitis findings survived FDR correction.

**Table 5.** Significant microstructural differences by performance status in the LGI1 encephalitis cohort.

| Navigation | DTI metric | Left hemisphere |  |  |  |  |  | Right hemisphere |  |  |  |  |  |
| --- | --- | --- | --- | --- | --- | --- | --- | --- | --- | --- | --- | --- | --- |
|  |  | n | < normal | controls | normal | g | p | n | < normal | controls | normal | g | p |
| Hippocampus | MD |  |  |  |  |  |  | 10 | 0.839 | 0.795 |  | 0.56 | .042 |
|  | FA |  |  |  |  |  |  | 11 | 0.112 | 0.120 |  | -0.53 | .044 |
| **Memory** |  |  |  |  |  |  |  |  |  |  |  |  |  |
| Hippocampus | MD | 11 | 0.903 | 0.798 |  | 0.77 | .010 | 10 | 0.845 | 0.799 |  | 0.57 | .041 |
|  | MD | 11/5 | 0.899 |  | 0.812 | 0.73 | .030 |  |  |  |  |  |  |
|  | FA | 11 | 0.117 | 0.126 |  | -0.53 | .042 |  |  |  |  |  |  |
| Cerebellar white matter | MD |  |  |  |  |  |  | 12/5 | 0.602 |  | 0.576 | 0.63 | .043 |
| Temporal cingulum | MD | 12 | 0.833 | 0.764 |  | 0.52 | .041 |  |  |  |  |  |  |
|  | MD | 12/5 | 0.830 |  | 0.763 | 0.83 | .029 |  |  |  |  |  |  |
|  | FA |  |  |  |  |  |  | 12 | 0.182 | 0.220 |  | -0.61 | .022 |

DTI = diffusion tensor imaging; FA = fractional anisotropy; MD = mean diffusivity; g = Hedges' g; MD values are reported in $10^{-3}$ mm<sup>2</sup>/s.

## Discussion

This study systematically examined spatial navigation and episodic memory function and their structural correlates in post-acute NMDAR and LGI1 encephalitis. Despite minimal residual disability, spatial navigation impairment was frequent, with convergent findings across two complementary paradigms and age as a common risk factor. In NMDAR encephalitis, spatial navigation emerged as a selectively vulnerable cognitive domain, with impairment occurring more frequently and largely independently of episodic memory impairment. This behavioural dissociation was mirrored by distinct neuroanatomical signatures, with navigation-specific alterations predominantly affecting paracentral and cerebellar regions, whereas memory-specific alterations more prominently involved the inferior temporal cortex, hippocampus and thalamus. In LGI1 encephalitis, navigation and memory impairments were more extensive and strongly overlapping. Structural alterations were more widespread and predominantly associated with memory impairment, involving lateral temporal and parietal cortices as well as the hippocampus and cerebellum, with no robust navigation-specific structural alterations. Together, these findings identify spatial navigation impairment as a frequent but under-recognized consequence of autoimmune encephalitis and reveal both distinct and overlapping neural signatures of navigation and memory impairment across its two most common subtypes.

Spatial navigation impairment emerged as a frequent sequela of both NMDAR and LGI1 encephalitis despite limited residual disability, affecting 57% and 70% of patients, respectively. These rates align with previous reports of spatial orientation complaints^6,35^ and visuoconstructive impairment in a nationwide Danish autoimmune encephalitis cohort^52^. They are also comparable to reported rates of persistent executive and working-memory impairment in NMDAR encephalitis up to five years after disease onset^5^. A nationwide Dutch study similarly found below-average performance in at least one cognitive domain in 65% of NMDAR encephalitis patients despite favourable functional outcomes, although basic visuoperceptual and constructional impairments were less frequent when using a stricter threshold^37^. In LGI1 encephalitis, earlier studies already identified persistent spatial disorientation even with comparatively coarse clinical assessments,^13,35^ consistent with the high frequency of navigation impairment and marked performance reductions observed here.

Importantly, episodic memory impairment in our NMDAR encephalitis cohort was at the lower end of prior prevalence estimates^5,8,10^. Although this may reflect the longer post-acute interval and exclusion of comorbid neurological conditions, it also shows the prominence of navigation impairment in this comparatively well-recovered cohort. Clinical reports of spatial disorientation and conventional visuospatial measures may therefore underestimate more subtle but persistent navigation difficulties. Given the heterogeneity of navigation tasks, convergent impairment across the complementary VIENNA Young and Sea Hero Quest paradigms strengthens the interpretation that identified impairment reflects an underlying navigation vulnerability rather than paradigm-specific effects^53^. Navigation impairment may therefore remain undetected when follow-up relies on orientation screening or standard memory tests, underscoring the need for a direct comparison with episodic memory function.

Behavioural findings in our study indicate that navigation dysfunction in autoimmune encephalitis cannot be understood solely as a downstream consequence of episodic memory impairment. In NMDAR encephalitis, selective navigation impairment was significantly more frequent than selective memory impairment. In LGI1 encephalitis, memory and navigation impairments overlapped strongly, with 17% of patients showing selective navigation impairment and 13% showing selective memory impairment. This relative predominance of navigation impairment remained when compared with visual memory and working memory, arguing against verbal test modality, general memory dysfunction or executive impairment as sufficient explanations. At the same time, working memory was the only cognitive domain associated with navigation performance, consistent with previous evidence and with the contribution of working memory to online spatial updating and maintenance of task-relevant spatial information^54–56^. This indicates that navigation draws on memory-related and executive processes but can nevertheless be selectively compromised even in the absence of evident impairment in either domain. This behavioural dissociation raises the question of whether navigation and memory dysfunction are also associated with separable neuroanatomical patterns.

In NMDAR encephalitis, the behavioural dissociation was mirrored by partly distinct, but not independent, macro- and microstructural alteration patterns. The strongest navigation-specific findings followed a parietal-cerebellar pattern, comprising bilateral paracentral and left parietal cortical thinning, reduced cerebellar grey matter volume and impaired cerebellar white matter integrity. This pattern aligns with evidence implicating paracentral and parietal cortices in object-location processing, egocentric perspective coding, reference-frame transformations, navigation performance in subjective cognitive decline and navigation-related cortical plasticity^32,57–59^.

The cerebellar findings are particularly notable in the context of established cerebellar vulnerability in NMDAR encephalitis^20,21^ and the cerebellum’s role in integrating idiothetic and visual information and coordinating with the hippocampus to support efficient navigation and stable spatial representations^34^. The separation between navigation- and memory-related patterns was nevertheless incomplete, for example with respect to shared hippocampal involvement: right hippocampal volume was reduced in both impairment groups, while hippocampal fractional anisotropy was associated specifically with navigation impairment and increased hippocampal mean diffusivity with memory impairment, consistent with prior navigation and NMDAR encephalitis memory findings^12,60–62^.

Cingulate and temporal white matter findings likewise indicated overlap: cingulate effects were distributed across navigation and memory analyses, and reduced temporal white matter integrity was associated with both navigation and memory, with more robust temporal cingulum involvement for memory. These findings are in line with evidence implicating disruption in cingulate and temporal pathways in NMDAR encephalitis and profound impact on memory- navigation networks^17,27,28,63–65^.

Memory-specific alterations more prominently involved left inferior temporal cortical thickness, left hippocampal volume and thalamic macro- and microstructure. The inferior temporal finding may partly reflect the verbal processing demands of the memory measures used here^66^, whereas the hippocampal and thalamic findings are consistent with evidence from neuroimmunological cohorts linking abnormalities in these structures to memory variability and cognitive impairment^11,67^. Overall, NMDAR findings support partial neuroanatomical differentiation, with the strongest navigation-specific effects in parietal and cerebellar regions, whereas memory impairment showed stronger temporal-hippocampal-thalamic involvement, alongside substantial overlap between domains for hippocampal, cingulate and temporal circuitry.

In LGI1 encephalitis, navigation and memory impairments were more extensive and substantially more overlapping, while structural alterations were primarily memory-related. This fits the LGI1 encephalitis cognitive phenotype, which is characterized by persistent executive and memory dysfunction^13,52,68,69^ and, structurally, by extensive temporal lobe involvement including hippocampal atrophy and cortical thinning associated with verbal memory dysfunction^9,13,18,70^. Although parietal, postcentral, hippocampal and cerebellar findings were also observed in association with navigation impairment, these alterations were more extensive and robustly associated with memory, indicating disruption of shared navigation-memory circuitry rather than a navigation-specific structural pattern^13,71^. Cingulate involvement was narrower than in NMDAR encephalitis, limited to navigation-related isthmus cortical thinning and memory-related temporal cingulum alterations, contributing to previous links between cingulate white matter alterations in LGI1 encephalitis and working and verbal memory^18,19^.

The absence of robust navigation-specific structural alterations suggests more global disruption of shared medial temporal and posterior cortical systems rather than a selective navigation-related vulnerability. Analogous evidence from ageing and Alzheimer’s disease suggests that navigation may reveal subtle network dysfunction early, whereas broader pathology produces increasingly severe and overlapping navigation and memory impairment^72–74^. The clinical separability of navigation and memory may therefore be greatest when network disruption is selective and become less apparent as pathology extends across shared neural systems.

Together, these complementary behavioural and neuroanatomical patterns within a shared clinical framework support a network-based view of spatial navigation and episodic memory as partially dissociable functions supported by overlapping and flexibly interacting hippocampal and extra-hippocampal systems^75^. Both disorders are neuronal-surface antibody-mediated encephalitides with prominent involvement of memory-related hippocampal-limbic networks, but they differ in target antigen and anatomical expression. Their comparison therefore suggests that navigation- memory separability depends on how pathology is distributed across shared and domain-weighted systems.

In line with relational memory accounts, navigation partly depends on hippocampal mechanisms for flexible relational organisation^76,77^. However, the navigation-specific contribution of the hippocampus may lie in the continuous, action-dependent integration of visual, idiothetic and task-dependent spatial information^78^. This demand is consistent with lesion evidence showing partial preservation of navigation despite medial temporal damage and even dense amnesia, particularly when multisensory input supports extrahippocampal compensation,^29,79,80^ and with meta-analytic evidence for navigation-related parietal and cerebellar involvement alongside memory-shared hippocampal and posterior cingulate contributions^81^.

These findings also have direct clinical implications, given the high frequency of below-normal navigation performance and the central role of spatial navigation for independence, safety and participation in daily life^24,82^. Yet navigation remains rarely assessed in routine clinical settings, partly because validated, reliable and norm-referenced measures have only recently become available^39,83–85^ and because episodic or visuospatial memory performance cannot serve as reliable proxies for navigation ability^38^. The brief and norm-referenced assessments used here could therefore complement longitudinal follow-up, particularly in younger or otherwise well-recovered patients whose difficulties may not be captured by coarse orientation items. Broader implications for selecting age-appropriate, psychometrically robust outcome measures for repeated neuropsychological follow-up are discussed in the Supplementary Discussion.

The age-related vulnerability observed here further suggests that navigation difficulties may become increasingly relevant during extended post-acute trajectories, when disease-related network alterations may exacerbate established age-related decline in navigation ability^86^. In NMDAR encephalitis, the association between longer time since onset and navigation impairment should not be interpreted as progressive worsening, which was not supported by longitudinal data, but more likely reflects historical improvements in recognition, diagnosis and immunotherapy^87^. These findings highlight the importance of including individually interpretable navigation measures in longitudinal outcome assessment. They also underscore the need to translate promising navigation training and support approaches into clinically applicable interventions with demonstrated efficacy across different levels of impairment^88–91^.

Some limitations should be considered. First, we used a relatively liberal threshold of PR ≤ 24 to identify below-normal performance. This prioritised sensitivity but should not be equated with functional impairment, as it includes both low-average and below-average performance. A stricter threshold was difficult to apply consistently because of the broad categorization bins provided by the German RAVLT norms^43^. We therefore used the lower bound of percentile ranges, which may modestly inflate episodic memory impairment rates when ranges crossed the PR 24 threshold; importantly, this would bias against, rather than favour, the detection of isolated navigation impairment. Second, premorbid ability was unknown, and the closest spatial-episodic comparator could not be analysed consistently because the established complex figure test^92^ showed marked ceiling effects and limited normative precision, whereas Brief Visuospatial Memory Test–Revised (BVMT-R)^93^ data were not available in sufficient numbers. Third, macro- and microstructural analyses were restricted to regions of primary interest because of the moderate sample sizes achievable in monocentre autoimmune encephalitis cohorts, leaving possible effects in other regions unexplored. These constraints warrant replication in larger, multicentre and multilingual cohorts, ideally combining longitudinal behavioural assessment with multimodal imaging. Such studies are feasible because both navigation paradigms are available in several languages and free for research use. Furthermore, identified performance levels should be linked to everyday outcomes such as participation, independent mobility and driving.

Taken together, our study identifies spatial navigation impairment as a frequent but under-assessed sequela of post-acute NMDAR and LGI1 encephalitis that cannot be inferred from episodic memory performance alone. By comparing two antibody-mediated encephalitis variants within a shared clinical and analytic framework, the study provides clinical evidence that navigation and episodic memory are partially dissociable but not independent. In NMDAR encephalitis, navigation and memory were more clearly differentiated at both behavioural and structural levels, whereas in LGI1 encephalitis they overlapped more strongly and were accompanied by widespread, predominantly memory-related structural alterations. These findings support a network-based account in which navigation-memory separability varies with the extent and distribution of pathology, and they argue for norm-referenced navigation assessment and targeted intervention strategies in longitudinal follow-up after autoimmune encephalitis.

## Data availability

All *R* scripts and data are openly available at the OSF under doi.org/10.17605/OSF.IO/B3FWZ and specify exact package versions used in the analyses. The navigation paradigms are available under doi.org/10.17605/OSF.IO/4H65P for VIENNA Young and celest.seaheroquest.com/ for Sea Hero Quest.

## Acknowledgements

We thank all participants and their families and supporters for their participation and commitment to this study. We are especially grateful to our research assistants, whose dedicated support in study coordination, data collection, and management was essential to the conduct of this study.

