## Supplementary Material for "Spatial navigation impairment beyond episodic memory in autoimmune encephalitis"

#### Supplementary Methods

#### Validation of normative model extrapolation to ages 68–84

To evaluate the extrapolation, we applied the normative model to healthy control participants aged 68 years or older. All observed scores fell within their corresponding 95% prediction intervals. This analysis cannot establish the validity of the extrapolation for individual-level interpretation, which would require a larger validation sample. However, the non-significant Kolmogorov–Smirnov test ($p$ = .092) and scatter of observed scores around the identity line provided no evidence against extrapolating the model across ages 68–84.

#### Supplementary Results

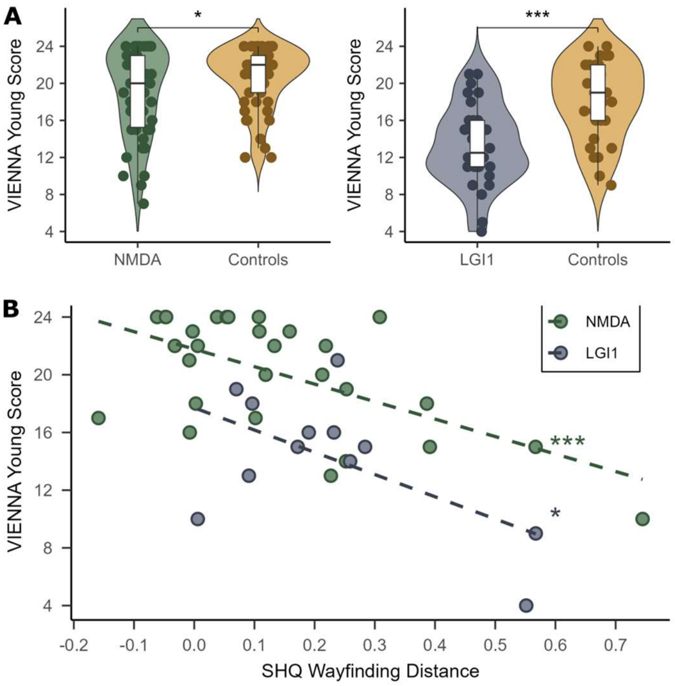


**Supplementary Figure 1. Navigation performance in patients with autoimmune encephalitis and age- and sex-matched control participants and convergence across navigation paradigms.** (**A**) Violin plots showing group differences in VIENNA Young performance between patients with NMDAR encephalitis and LGI1 encephalitis and their respective age- and sex-matched controls. (**B**) Scatterplot and linear regression lines showing the association between the two navigation measures VIENNA Young and Sea Hero Quest (SHQ) mean wayfinding distance, normalized to distance travelled during practice levels 1 and 2 (dexterity-adjusted).

#### Visual episodic and working memory validation analyses

As a validation analysis, we tested whether the pattern of relatively greater navigation than memory impairment in NMDAR encephalitis could be explained by differences in verbal versus visual test modality. To do so, we derived a binary visual memory score in a subset of 49 NMDAR and 28 LGI1 encephalitis patients for whom visual memory data were available. Owing to a protocol change during data collection, this score was based on different instruments across patients: in the NMDAR encephalitis group, visual memory was derived from the delayed recall trial of the Rey–Osterrieth Complex Figure Test (ROCF) (Rey, 1941) in 32 patients and from the sum of learning trials and delayed recall of the Brief Visuospatial Memory Test–Revised (BVMT-R) (Langdon et al., 2012) in the remaining 17 patients. In the LGI1 encephalitis cohort, visual memory scores were derived from the ROCF in 12 patients and from the BVMT-R in 16 patients.

Consistent with the findings for verbal episodic memory, isolated spatial navigation impairment was significantly more prevalent than isolated visual episodic memory impairment in the NMDAR encephalitis group ($\chi^2(1) = 15.70$, $p < .001$) and also in the LGI1 encephalitis group (exact McNemar's test, $p = .007$; Supplementary Fig. 2A and C).

We next compared navigation impairment with working memory impairment, assessed with digit span backwards, as another cognitive measure known to be particularly vulnerable in NMDAR encephalitis and relevant to executive control. Again, isolated spatial navigation impairment was significantly more frequent than isolated working memory impairment in both the NMDAR encephalitis group ($\chi^2(1) = 11.64$, $p < .001$) and the LGI1 encephalitis group (exact McNemar's test, $p < .001$; Supplementary Fig. 2B and D).

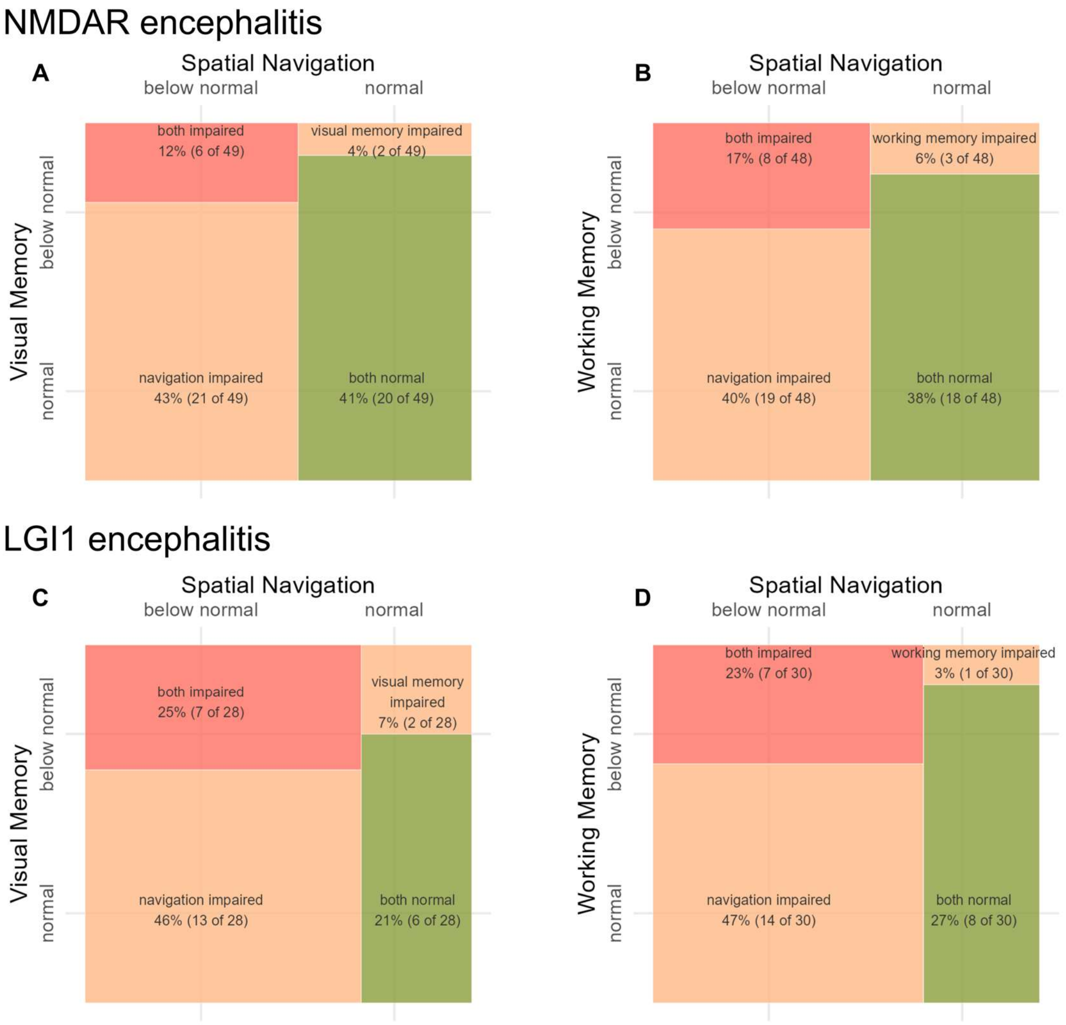


**Supplementary Figure 2. Mosaic plots of binary spatial navigation, visual memory, and working memory status in NMDAR encephalitis and LGI1 encephalitis**. Mosaic plots showing the overlap between spatial navigation and visual episodic memory (**A**, **C**) and between spatial navigation and working memory (**B**, **D**) in NMDAR encephalitis (**A**, **B**) and LGI1 encephalitis (**C**, **D**). Spatial navigation was based on both navigation markers (VIENNA Young and SHQ). Visual memory was based on at least one visual episodic memory measure (ROCF or BVMT-R), and working memory was assessed with digit span backwards. All measures were categorized as within normal expectations (PR > 24) or below normal expectations (PR ≤ 24).

#### Cognitive correlates of navigation using continuous test scores

To reduce potential bias arising from differences between the normative references used across cognitive tests, we conducted separate multiple regression analyses using continuous test performance. VIENNA Young performance was entered as the dependent variable, with working memory, delayed recall, and attention, measured by median reaction time in a tonic alertness task, entered as cognitive predictors. All models were adjusted for age, sex, and education.

In NMDAR encephalitis, working memory was significantly associated with navigation performance ($b = 1.00$, $t(40) = 3.56$, $p < .001$), whereas delayed recall and attention were not significantly associated with navigation. The model explained 50% of the variance in VIENNA Young performance. In LGI1 encephalitis, none of the cognitive predictors were significantly associated with navigation performance, although the full model explained 55% of the variance.

### General macrostructural group comparisons

##### Table S1 Macrostructural group comparisons in the NMDAR encephalitis cohort.

| Region | Left hemisphere | | | | Right hemisphere | | | |
|---|---|---|---|---|---|---|---|---|
| | Patients *M* (*SD*) | Controls *M* (*SD*) | *g* | *p* | Patients *M* (*SD*) | Controls *M* (*SD*) | *g* | *p* |
| **Cortical thickness (mm)** | | | | | | | | |
| Entorhinal | 3.17 (0.25) | 3.24 (0.25) | -0.18 | .121 | 3.35 (0.26) | 3.31 (0.25) | 0.08 | .705 |
| Parahippocampal | 2.76 (0.27) | 2.74 (0.22) | 0.05 | .640 | 2.69 (0.25) | 2.68 (0.22) | 0.00 | .512 |
| Fusiform | 2.71 (0.12) | 2.73 (0.11) | -0.07 | .311 | 2.69 (0.14) | 2.71 (0.11) | -0.12 | .210 |
| Superior temporal | 2.79 (0.18) | 2.87 (0.13) | -0.35 | .010 | 2.79 (0.16) | 2.85 (0.15) | -0.31 | .019 |
| Middle temporal | 2.82 (0.15) | 2.87 (0.13) | -0.30 | .024 | 2.82 (0.14) | 2.86 (0.14) | -0.21 | .079 |
| Inferior temporal | 2.76 (0.14) | 2.80 (0.11) | -0.22 | .072 | 2.76 (0.12) | 2.75 (0.12) | 0.05 | .632 |
| Transverse temporal | 2.41 (0.22) | 2.40 (0.20) | 0.04 | .600 | 2.42 (0.26) | 2.42 (0.18) | -0.00 | .494 |
| Banks STS | 2.44 (0.17) | 2.49 (0.19) | -0.17 | .123 | 2.58 (0.18) | 2.63 (0.20) | -0.19 | .095 |
| Paracentral | 2.41 (0.13) | 2.47 (0.14) | -0.32 | .014 | 2.48 (0.16) | 2.50 (0.15) | -0.09 | .264 |
| Postcentral | 2.11 (0.11) | 2.14 (0.12) | -0.23 | .058 | 2.11 (0.14) | 2.13 (0.14) | -0.12 | .209 |
| Supramarginal | 2.59 (0.13) | 2.62 (0.12) | -0.20 | .086 | 2.59 (0.13) | 2.62 (0.11) | -0.21 | .073 |
| Superior parietal | 2.19 (0.10) | 2.22 (0.10) | -0.23 | .058 | 2.18 (0.11) | 2.21 (0.10) | -0.21 | .072 |
| Inferior parietal | 2.48 (0.11) | 2.49 (0.10) | -0.04 | .392 | 2.52 (0.11) | 2.56 (0.10) | -0.31 | .016 |
| Precuneus | 2.38 (0.10) | 2.40 (0.10) | -0.07 | .322 | 2.41 (0.12) | 2.42 (0.10) | -0.05 | .367 |
| Isthmus-cingulate | 2.31 (0.17) | 2.34 (0.18) | -0.10 | .235 | 2.31 (0.17) | 2.34 (0.11) | -0.16 | .127 |
| Posterior-cingulate | 2.41 (0.17) | 2.44 (0.14) | -0.15 | .152 | 2.38 (0.16) | 2.42 (0.12) | -0.20 | .084 |
| Caudal anterior cingulate | 2.49 (0.24) | 2.59 (0.16) | -0.33 | .012 | 2.33 (0.21) | 2.38 (0.17) | -0.18 | .107 |
| Rostral anterior cingulate | 2.75 (0.19) | 2.81 (0.14) | -0.23 | .053 | 2.63 (0.19) | 2.66 (0.17) | -0.15 | .151 |
| **eTIV adjusted volume (mm³)** | | | | | | | | |
| Hippocampus | 4,033 (446) | 4,290 (391) | -0.27 | .056 | 4,157 (432) | 4,446 (451) | -0.34* | .021 |
| Thalamus | 7,420 (991) | 7,756 (812) | -0.03 | .402 | 7,312 (856) | 7,519 (654) | 0.11 | .836 |
| Cerebellum | 52,940 (5,793) | 57,221 (4,982) | -0.31* | .005 | 53,489 (6,189) | 57,961 (5,353) | -0.30* | .006 |

*M* = mean; *SD* = standard deviation; *$p_{FDR}$ false discovery rate corrected *p*-value < 0.05.

##### Table S2 Macrostructural group comparisons in the LGI1 encephalitis cohort.

| Region | Left hemisphere | | | | Right hemisphere | | | |
|---|---|---|---|---|---|---|---|---|
| | Patients *M* (*SD*) | Controls *M* (*SD*) | *g* | *p* | Patients *M* (*SD*) | Controls *M* (*SD*) | *g* | *p* |
| **Cortical thickness (mm)** | | | | | | | | |
| Entorhinal | 3.08 (0.38) | 3.28 (0.27) | -0.42* | .014 | 3.17 (0.36) | 3.29 (0.21) | -0.25 | .089 |
| Parahippocampal | 2.60 (0.31) | 2.69 (0.29) | -0.18 | .158 | 2.55 (0.30) | 2.63 (0.21) | -0.18 | .165 |
| Fusiform | 2.59 (0.14) | 2.68 (0.13) | -0.49* | .006 | 2.57 (0.16) | 2.66 (0.14) | -0.42* | .013 |
| Superior temporal | 2.65 (0.18) | 2.68 (0.19) | -0.19 | .154 | 2.62 (0.19) | 2.71 (0.19) | -0.36 | .029 |
| Middle temporal | 2.68 (0.14) | 2.78 (0.18) | -0.59* | .001 | 2.69 (0.17) | 2.77 (0.16) | -0.30 | .055 |
| Inferior temporal | 2.63 (0.13) | 2.72 (0.16) | -0.41* | .016 | 2.64 (0.13) | 2.72 (0.14) | -0.44* | .011 |
| Transverse temporal | 2.15 (0.28) | 2.26 (0.22) | -0.29 | .061 | 2.20 (0.30) | 2.29 (0.28) | -0.24 | .093 |
| Banks STS | 2.31 (0.16) | 2.38 (0.16) | -0.37 | .026 | 2.42 (0.18) | 2.50 (0.19) | -0.33 | .041 |
| Paracentral | 2.29 (0.16) | 2.37 (0.14) | -0.36 | .028 | 2.32 (0.20) | 2.42 (0.15) | -0.36 | .029 |
| Postcentral | 1.97 (0.15) | 2.01 (0.14) | -0.24 | .095 | 1.94 (0.17) | 2.01 (0.14) | -0.35 | .032 |
| Supramarginal | 2.38 (0.16) | 2.48 (0.12) | -0.53* | .003 | 2.35 (0.18) | 2.49 (0.16) | -0.59* | .001 |
| Superior parietal | 2.08 (0.14) | 2.14 (0.10) | -0.32 | .043 | 2.04 (0.15) | 2.11 (0.14) | -0.35 | .031 |
| Inferior parietal | 2.31 (0.14) | 2.38 (0.12) | -0.45* | .010 | 2.34 (0.16) | 2.41 (0.14) | -0.36 | .030 |
| Precuneus | 2.20 (0.16) | 2.30 (0.13) | -0.48* | .007 | 2.21 (0.19) | 2.32 (0.13) | -0.48* | .007 |
| Isthmus-cingulate | 2.18 (0.13) | 2.29 (0.16) | -0.52* | .004 | 2.18 (0.19) | 2.27 (0.17) | -0.41* | .016 |
| Posterior-cingulate | 2.35 (0.10) | 2.42 (0.18) | -0.26 | .087 | 2.29 (0.13) | 2.38 (0.17) | -0.42* | .013 |
| Caudal anterior cingulate | 2.44 (0.24) | 2.50 (0.25) | -0.16 | .197 | 2.33 (0.18) | 2.34 (0.21) | -0.07 | .358 |
| Rostral anterior cingulate | 2.63 (0.25) | 2.69 (0.20) | -0.18 | .160 | 2.63 (0.21) | 2.59 (0.23) | 0.11 | .720 |
| **eTIV adjusted volume (mm³)** | | | | | | | | |
| Hippocampus | 3,854 (715) | 4,242 (510) | -0.35 | .033 | 4,065 (698) | 4,406 (522) | -0.31 | .033 |
| Thalamus | 7,184 (998) | 7,397 (1,090) | -0.01 | .464 | 7,197 (918) | 7,364 (970) | 0.00 | .502 |
| Cerebellum | 52,816 (5,585) | 56,425 (7,547) | -0.26 | .078 | 54,548 (5,760) | 57,621 (7,944) | -0.17 | .171 |

*M* = mean; *SD* = standard deviation; *$p_{FDR}$ false discovery rate corrected *p*-value < 0.05.

### General microstructural group comparisons

##### Table S3 Microstructural group comparisons in the NMDAR encephalitis cohort.

| | | Left hemisphere | | | | | Right hemisphere | | | | |
|---|---|---|---|---|---|---|---|---|---|---|---|
| Region | DTI metric | n pairs | Patients *M* (*SD*) | Controls *M* (*SD*) | *g* | *p* | n pairs | Patients *M* (*SD*) | Controls *M* (*SD*) | *g* | *p* |
| Hippocampus | MD | 41 | 0.798 (0.035) | 0.785 (0.017) | 0.29 | .033 | 42 | 0.786 (0.026) | 0.784 (0.019) | 0.06 | .358 |
| | FA | 41 | 0.122 (0.015) | 0.131 (0.011) | -0.57** | < .001 | 41 | 0.124 (0.014) | 0.131 (0.014) | -0.42* | .004 |
| Thalamus | MD | 43 | 0.661 (0.018) | 0.664 (0.013) | -0.12 | .790 | 42 | 0.663 (0.018) | 0.661 (0.011) | 0.08 | .301 |
| | FA | 43 | 0.285 (0.011) | 0.289 (0.014) | -0.25 | .051 | 43 | 0.282 (0.014) | 0.284 (0.011) | -0.08 | .291 |
| Cerebellar white matter | MD | 43 | 0.603 (0.018) | 0.592 (0.014) | 0.50** | < .001 | 42 | 0.603 (0.018) | 0.590 (0.014) | 0.56** | < .001 |
| | FA | 43 | 0.455 (0.031) | 0.465 (0.024) | -0.35 | .013 | 42 | 0.457 (0.031) | 0.467 (0.025) | -0.32 | .021 |
| Dorsal cingulum | MD | 37 | 0.678 (0.018) | 0.674 (0.021) | 0.13 | .216 | 37 | 0.704 (0.130) | 0.680 (0.019) | 0.18 | .135 |
| | FA | 37 | 0.379 (0.033) | 0.385 (0.034) | -0.13 | .210 | 37 | 0.355 (0.056) | 0.363 (0.033) | -0.11 | .259 |
| Peri-genual cingulum | MD | 31 | 0.713 (0.022) | 0.703 (0.036) | 0.23 | .097 | 32 | 0.716 (0.024) | 0.721 (0.048) | -0.10 | .716 |
| | FA | 30 | 0.338 (0.041) | 0.352 (0.050) | -0.27 | .067 | 32 | 0.309 (0.045) | 0.325 (0.051) | -0.25 | .079 |
| Temporal cingulum | MD | 39 | 0.748 (0.029) | 0.739 (0.021) | 0.27 | .049 | 39 | 0.749 (0.029) | 0.739 (0.024) | 0.30 | .030 |
| | FA | 39 | 0.226 (0.044) | 0.221 (0.037) | 0.09 | .705 | 41 | 0.229 (0.042) | 0.236 (0.035) | -0.17 | .141 |
| Fornix | MD | 8 | 0.715 (0.034) | 0.744 (0.022) | -0.57 | .943 | 6 | 0.795 (0.065) | 0.769 (0.062) | 0.28 | .226 |
| | FA | 8 | 0.319 (0.043) | 0.287 (0.070) | 0.29 | .808 | 6 | 0.250 (0.046) | 0.269 (0.059) | -0.26 | .238 |

FA = fractional anisotropy; MD = mean diffusivity; *g* = Hedges' *g*; MD values are reported in $10^{-3}$ $mm^2/s$; *$p_{FDR}$ false discovery rate corrected *p*-value < 0.05; **$p_{FDR}$ false discovery rate corrected *p*-value < 0.01.

##### Table S4 Microstructural group comparisons in the LGI1 encephalitis cohort.

| | | Left hemisphere | | | | | Right hemisphere | | | | |
|---|---|---|---|---|---|---|---|---|---|---|---|
| Region | DTI metric | n pairs | Patients *M* (*SD*) | Controls *M* (*SD*) | *g* | *p* | n pairs | Patients *M* (*SD*) | Controls *M* (*SD*) | *g* | *p* |
| Hippocampus | MD | 16 | 0.872 (0.117) | 0.795 (0.020) | 0.65 | .008 | 15 | 0.830 (0.064) | 0.793 (0.018) | 0.51 | .029 |
| | FA | 16 | 0.116 (0.011) | 0.126 (0.019) | -0.50 | .026 | 16 | 0.113 (0.007) | 0.119 (0.012) | -0.52 | .022 |
| Thalamus | MD | 16 | 0.667 (0.017) | 0.669 (0.023) | -0.06 | .595 | 16 | 0.674 (0.017) | 0.670 (0.022) | 0.16 | .255 |
| | FA | 16 | 0.301 (0.017) | 0.295 (0.012) | 0.38 | .933 | 16 | 0.292 (0.014) | 0.287 (0.012) | 0.43 | .954 |
| Cerebellar white matter | MD | 15 | 0.598 (0.042) | 0.597 (0.013) | 0.02 | .475 | 17 | 0.594 (0.039) | 0.592 (0.016) | 0.05 | .415 |
| | FA | 15 | 0.454 (0.040) | 0.463 (0.029) | -0.24 | .175 | 16 | 0.465 (0.043) | 0.467 (0.032) | -0.05 | .424 |
| Dorsal cingulum | MD | 15 | 0.689 (0.029) | 0.685 (0.024) | 0.11 | .335 | 15 | 0.696 (0.029) | 0.694 (0.026) | 0.03 | .449 |
| | FA | 15 | 0.360 (0.031) | 0.362 (0.028) | -0.06 | .402 | 15 | 0.336 (0.027) | 0.334 (0.030) | 0.06 | .590 |
| Peri-genual cingulum | MD | 16 | 0.719 (0.039) | 0.709 (0.031) | 0.17 | .238 | 15 | 0.727 (0.033) | 0.723 (0.028) | 0.07 | .396 |
| | FA | 15 | 0.308 (0.033) | 0.325 (0.035) | -0.34 | .095 | 15 | 0.287 (0.035) | 0.301 (0.033) | -0.30 | .117 |
| Temporal cingulum | MD | 17 | 0.811 (0.089) | 0.757 (0.044) | 0.47 | .029 | 16 | 0.781 (0.044) | 0.761 (0.044) | 0.36 | .076 |
| | FA | 17 | 0.194 (0.030) | 0.210 (0.033) | -0.35 | .077 | 17 | 0.185 (0.031) | 0.217 (0.039) | -0.58 | .012 |
| Fornix | MD | 7 | 0.791 (0.070) | 0.743 (0.065) | 0.44 | .114 | 8 | 0.913 (0.287) | 0.756 (0.055) | 0.48 | .086 |
| | FA | 7 | 0.257 (0.040) | 0.312 (0.085) | -0.61 | .056 | 8 | 0.269 (0.067) | 0.310 (0.067) | -0.39 | .127 |

FA = fractional anisotropy; MD = mean diffusivity; *g* = Hedges' *g*; MD values are reported in $10^{-3}$ $mm^2/s$; *$p_{FDR}$ false discovery rate corrected *p*-value < 0.05; **$p_{FDR}$ false discovery rate corrected *p*-value < 0.01.

### Supplementary Discussion

More broadly, neuropsychological follow-up after autoimmune encephalitis should prioritise age-appropriate, clinically meaningful outcomes that allow individual change to be interpreted over time. This is particularly important when premorbid ability is unknown and recovery may differ across domains, with memory improving while visual-spatial or executive deficits persist. Measures should therefore be selected not only for tradition, face validity or similarity to everyday behaviour, but for demonstrated criterion and predictive validity, adequate reliability, normative referencing and suitability for repeated assessment (Suchy et al., 2024). Virtual or everyday-like paradigms may increase face validity, but they require the same psychometric standards as conventional tests. In younger or well-recovered patients, established memory measures may be insufficiently sensitive to residual impairment and longitudinal change because of ceiling effects, broad percentile bins, outdated normative data and familiarity effects with repeated administration, particularly in complex figure paradigms. Follow-up should therefore combine conventional cognitive testing with age-appropriate navigation and visual-spatial measures and, where possible, use reliable-change indices or comparable individual-level change metrics rather than static impairment thresholds alone. Measures with low quality and especially outdated normative foundations, limited sensitivity to change or little relevance to clinical management may warrant lower priority in repeated assessments.
